# Conservative and liberal attitudes drive polarized neural responses to political content

**DOI:** 10.1101/2020.04.30.071084

**Authors:** Yuan Chang Leong, Janice Chen, Robb Willer, Jamil Zaki

## Abstract

People tend to interpret political information in a manner that confirms their prior beliefs, a cognitive bias that contributes to rising political polarization. In this study, we combined functional magnetic resonance imaging with semantic content analyses to investigate the neural mechanisms that underlie the biased processing of real-world political content. We scanned American participants with conservative-leaning or liberal-leaning immigration attitudes while they watched news clips, campaign ads, and public speeches related to immigration policy. We searched for evidence of “neural polarization”: activity in the brain that diverges between people who hold liberal versus conservative political attitudes. Neural polarization was observed in the dorsomedial prefrontal cortex (DMPFC), a brain region associated with the interpretation of narrative content. Neural polarization in the DMPFC intensified during moments in the videos that included risk-related and moral-emotional language, highlighting content features most likely to drive divergent interpretations between conservatives and liberals. Finally, participants whose DMPFC activity closely matched that of the average conservative or the average liberal participant were more likely to change their attitudes in the direction of that group’s position. Our work introduces a novel multi-method approach to study the neural basis of political cognition in naturalistic settings. Using this approach, we characterize how political attitudes biased information processing in the brain, the language most likely to drive polarized neural responses, and the consequences of biased processing for attitude change. Together, these results shed light on the psychological and neural underpinnings of how identical information is interpreted differently by conservatives and liberals.

**Significance Statement:** Partisan biases in processing political information contribute to rising divisions in society. How do such biases arise in the brain? We measured the neural activity of participants watching videos related to immigration policy. Despite watching the same videos, conservative and liberal participants exhibited divergent neural responses. This “neural polarization” between groups occurred in a brain area associated with the interpretation of narrative content, and intensified in response to language associated with risk, emotion, and morality. Furthermore, polarized neural responses predicted attitude change in response to the videos. These findings suggest that biased processing in the brain drives divergent interpretations of political information and subsequent attitude polarization.

---

Political polarization is a growing concern in societies across the world (1). In the US, Democrats and Republicans have grown more ideologically divided in recent years, threatening both social harmony and effective governance (2, 3). Motivated political reasoning is a robust phenomenon thought to contribute to political polarization (4, 5). When presented with identical information, individuals with opposing political attitudes often become more entrenched in their original positions (6–9). The biased assimilation of political information impedes efforts to persuade partisans towards positions of consensus and compromise.

Why does the same information trigger divergent responses across individuals? One possibility is that motivation biases sensory attention (10, 11), such that people attend more to information that supports their beliefs. For example, when watching news footage about a protest, detractors of the protest might focus on aspects of the video suggesting that protestors are behaving in a threatening manner so as to discredit their cause. Consistent with this view, previous work suggests that political attitudes bias processing as early as sensory perception (12, 13). Alternatively, motivation might affect how people interpret the same sensory input. That is, the same actions can be interpreted as threatening or not threatening depending on one’s prior attitudes.

Neuroscience can offer new insights into the fundamental cognitive processes that give rise to motivated political reasoning (14, 15). By assessing when biases in information processing emerge in the brain (e.g., “early” sensory cortices versus “late” association cortices), we can better understand how political attitudes affect different levels of information processing. However, real-world political content (e.g., news clips, televised debates) is often dynamic and complex, and thus incompatible with typical neuroscientific analytical approaches that require averaging over short, repeatable “trials”. This presents a challenge to researchers studying the neural mechanisms underlying the biased processing of political information. To our knowledge, no study has examined how the brain processes naturalistic audio-visual political content.

In the current study, we draw on advances in the analysis of functional magnetic resonance imaging (fMRI) data to examine how political attitudes bias the processing of naturalistic political content. We scanned conservative or liberal-leaning participants while they watched videos related to immigration policy, a polarized and politically significant topic in the U.S. (16) and in many countries around the world (17). Our analytical approach relies on the method of inter-subject correlation (ISC), which computes the correlation in activity between brains as a measure of shared processing. ISC has been previously used to examine how the brain processes naturalistic stimuli such as spoken narratives or films, and is well-suited to the study of the processing of real-world political content (18).

The first goal of our study is to examine if and how political attitudes modulate neural responses at different levels of information processing in the brain. Using ISC, we searched for evidence of *neural polarization* - activity that is shared between individuals with similar political attitudes but not between individuals with dissimilar political attitudes. Neural polarization measures the extent to which processing in a particular brain area diverges between conservative and liberal-leaning participants. If political attitudes bias sensory attention, we would expect neural polarization to emerge early in the processing stream (e.g., primary visual or auditory cortices) (19) Alternatively, if political attitudes bias the interpretation of the videos without altering sensory processing, we would expect neural polarization to emerge only in “higher-order” brain areas such as the dorsomedial prefrontal cortex (DMPFC), posterior medial cortex (PMC) or middle temporal gyrus (MTG). These brain regions have been previously shown to track the interpretation of narrative content (20, 21).

The second goal of our study was to characterize the content features in political information that were most likely to drive neural polarization. The literature on political psychology suggests that differences in political attitudes are associated with differences in moral values (22–24), and that emotional content enhances the polarizing effects of political messages (25, 26). We thus hypothesize that moral and emotional content would be viewed differently by participants with different political attitudes and thus most likely to drive polarized neural responses. To test this hypothesis, we first examined if moral and emotional content in the videos was associated with greater neural polarization. In a second analysis, we took advantage of the richness and complexity of our videos to test what other content features were likely to drive neural polarization. We broke down the content of the videos into 50 semantic categories (e.g., words related to risk, social affiliation and religion), and assessed the extent to which each category was associated with greater neural polarization. This allowed us to take a data-driven approach to identify content that contributed to polarized neural responses.

The third goal of our study was to examine the relationship between polarized neural responses and attitude change. If shared neural responses reflect shared interpretation of a video, we would expect the degree of neural similarity to be associated with attitude change after viewing the video. In particular, the degree to which a participant’s neural response was similar to that of conservative or liberal participants would predict attitude change towards more conservative or liberal positions on immigration, respectively. We tested this hypothesis by assessing if neural similarity to the average conservative or average liberal participant while watching each video would be associated with self-reported ratings of how much the video changed participants’ attitudes on the relevant immigration policy.

Our study introduces a new approach for investigating the neural processes underlying political cognition. In adapting fMRI paradigms to use real-world naturalistic political content, we study the biased processing of political content in a setting where we can be more confident of ecological validity. Using this approach, we identified a novel neural signature of biased processing of political information. The richness of naturalistic stimuli also allowed for data-driven analyses that generate new hypotheses to be examined in future experiments. By examining how the brain processes political content, our study advances our understanding of the partisan brain and how it gives rise to the polarization afflicting societies today.

## Results

Thirty-eight participants were scanned using fMRI as they watched twenty-four videos on six immigration policies (total duration = 35min 26s, divided into four runs; Fig. 1A, 1B). An online pre-test with a larger sample indicated that conservatives and liberals in America held opposing attitudes on these policies (*Supplemental Results*, Fig. S1). Prior to the experiment, participants indicated their support for each of the six policies. Their responses were recoded such that lower ratings reflect stronger support for liberal positions while higher ratings reflect stronger support for conservative positions. We then tallied each participant’s response to compute an “Immigration Attitude” score, and performed a median split to identify participants with conservative-leaning immigration attitudes and participants with liberal-leaning immigration attitudes (Fig. 1C). The two groups did not differ significantly on age, sex, income, education and amount of head motion in the scanner (Table S1).

**Figure 1.**
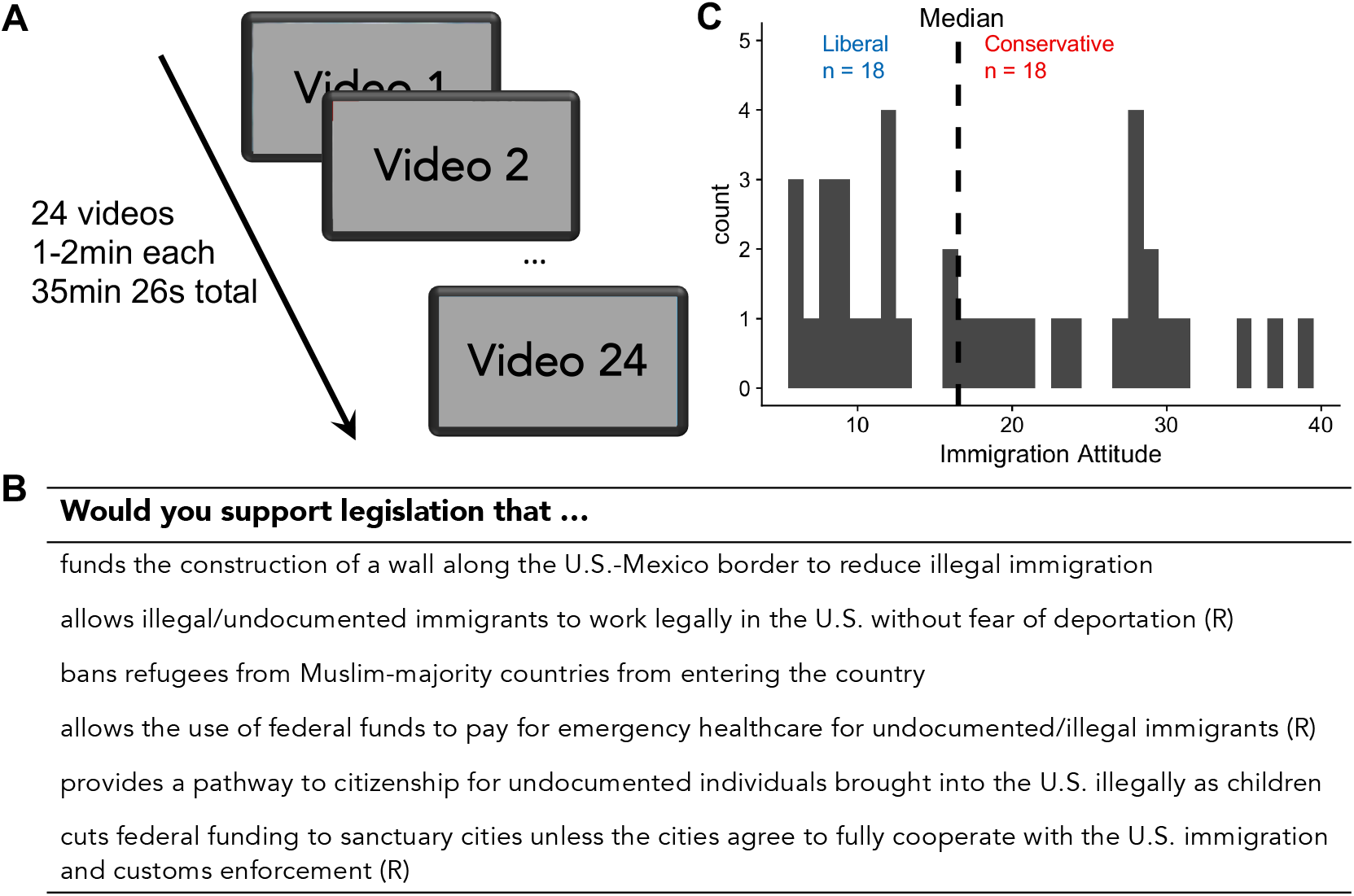
Experimental Design. **A.** Participants watched 24 videos on 6 immigration policies while undergoing fMRI. **B.** Prior to the experiment, participants indicated their support for each policy on a 7-point scale. Three of the questions, indicated here with (R), were reverse coded such that higher ratings on all questions indicate stronger support for the conservative position while lower ratings indicate stronger support for the liberal position. **C.** We tallied participants’ responses to compute their Immigration Attitude score, and performed a median split to identify liberal-leaning and conservative-leaning participants.

### Political content elicits shared neural responses across participants

We first examined the extent to which viewing political videos elicited similar neural responses across participants. For each participant, we z-scored the activity time course for each video and concatenated the neural data such that the videos were ordered in the same sequence. For each voxel in a participant’s brain, we calculated the intersubject-correlation (ISC) as the correlation between that voxel’s timecourse of activity and the average timecourse of all other participants at the same voxel (18). This correlation was computed across the entire duration of the twenty-four videos. The resulting *r* values were then averaged across participants to obtain a map of average *r* values, which shows the extent to which activity at a given voxel was similar across participants. Statistical significance was assessed using a permutation procedure where the sign of each individual participant’s ISC values was randomly flipped to generate a null distribution for each voxel (see Methods).

Consistent with earlier studies using audio-visual stimuli (27, 28), we observed high ISC in the primary auditory and visual cortices, and low ISC in motor and somatosensory cortices (Fig. 2). Similar to these studies, we also found widespread ISC in the medial and lateral prefrontal cortices (MPFC, LPFC), posterior medial cortex (PMC) and middle temporal gyrus (MTG), areas which have been previously associated with the processing and comprehension of narrative stimuli (20, 21). These results indicate that the political videos evoked reliable neural responses that are shared across participants, irrespective of their political attitudes.

**Figure 2.**
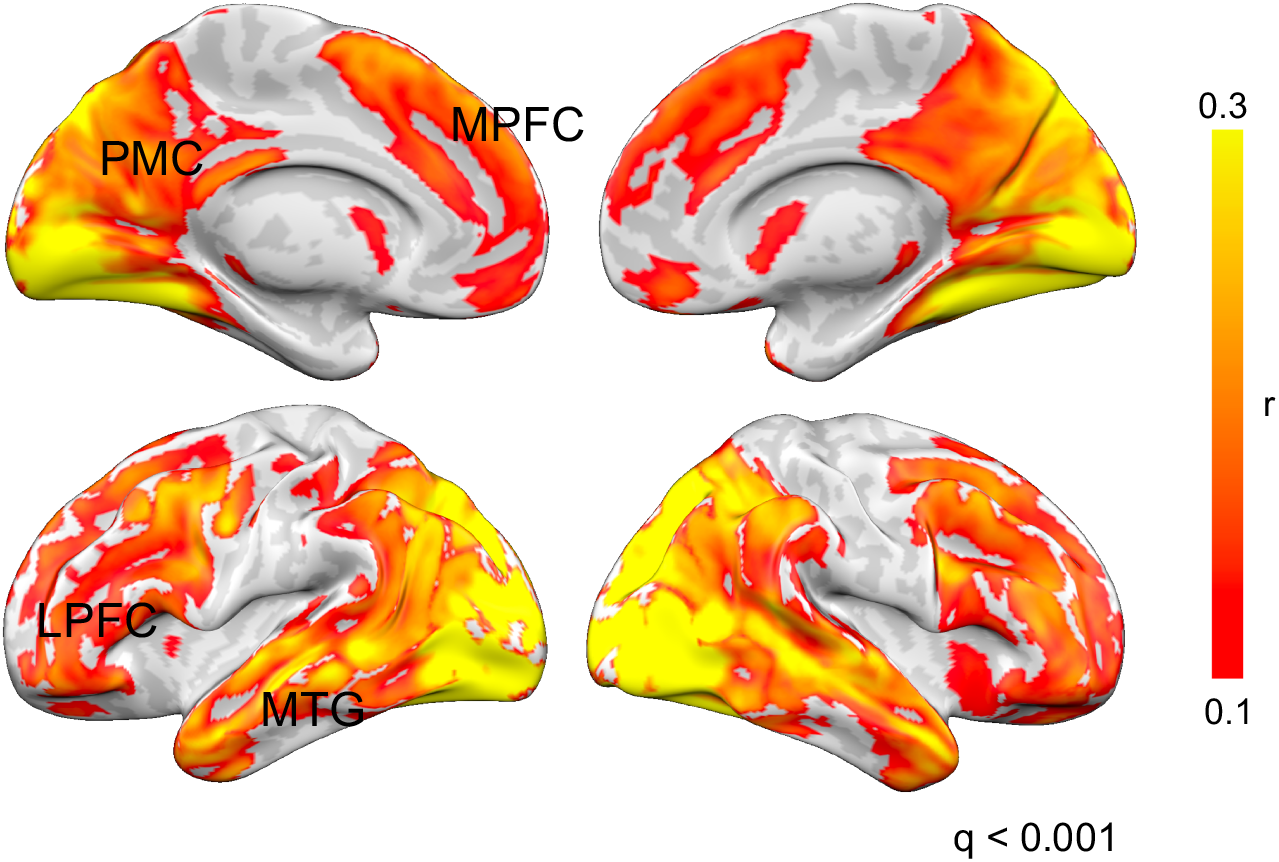
Shared neural response elicited by videos across participants. The videos elicited reliable shared responses across participants irrespective of their political attitudes. Intersubject correlation (ISC) was highest in sensory regions, including the primary visual and auditory cortices. There was also moderate ISC in higher-order regions such as the medial and lateral prefrontal cortices (MPFC, LPFC), posterior medial cortex (PMC) and middle temporal gyrus (MTG). Statistical maps were thresholded at a false discovery rate (FDR) of q < 0.001. Unthresholded map available at: https://neurovault.org/collections/PKFXOYLX/images/319401/

### DMPFC activity was more similar between participants with similar political attitudes

Our next analysis focused on identifying brain areas that exhibited evidence of neural polarization. For each participant, we computed a “within-group ISC” as their voxelwise ISC with the average timecourse of all other participants with similar political attitudes (i.e. liberal vs. average liberal and conservative vs. average conservative), and a “between-group ISC” as the ISC with the average timecourse of all participants with dissimilar political attitudes (i.e. liberal vs. average conservative; conservative vs. average liberal). The difference between within-group and between-group ISC measures the degree to which neural activity was shared between participants with similar political attitudes but not between participants with dissimilar political attitudes. We thus searched the brain for voxels where within-group ISC was greater than between-group ISC to identify brain areas where the processing diverged between the two groups.

If differences between groups were due to discrepancies in low-level visual or auditory attention, we would expect differences in ISC to emerge early in the processing stream (e.g., primary visual or auditory cortex). In contrast, if the differences were related to interpretation and evaluation of the same audio-visual input, we would expect to see differences emerge in “higher-order” association cortex. Consistent with the latter hypothesis, within-group ISC was greater than between-group ISC only in the left dorsomedial prefrontal cortex (DMPFC; Fig. 3A). Previous work using ambiguous stimuli have found that activity in this region tracks the interpretation of narratives, and is more similar between participants with similar interpretations of the same narrative (20, 21, 29). Within-group ISC in the DMPFC was higher than between-group ISC in both conservative and liberal participants, indicating that the results were not driven by one of the two groups (Fig. S2). In contrast, within-group ISC was not different from between-group ISC in primary sensory regions (Fig. 3B).

**Figure 3.**
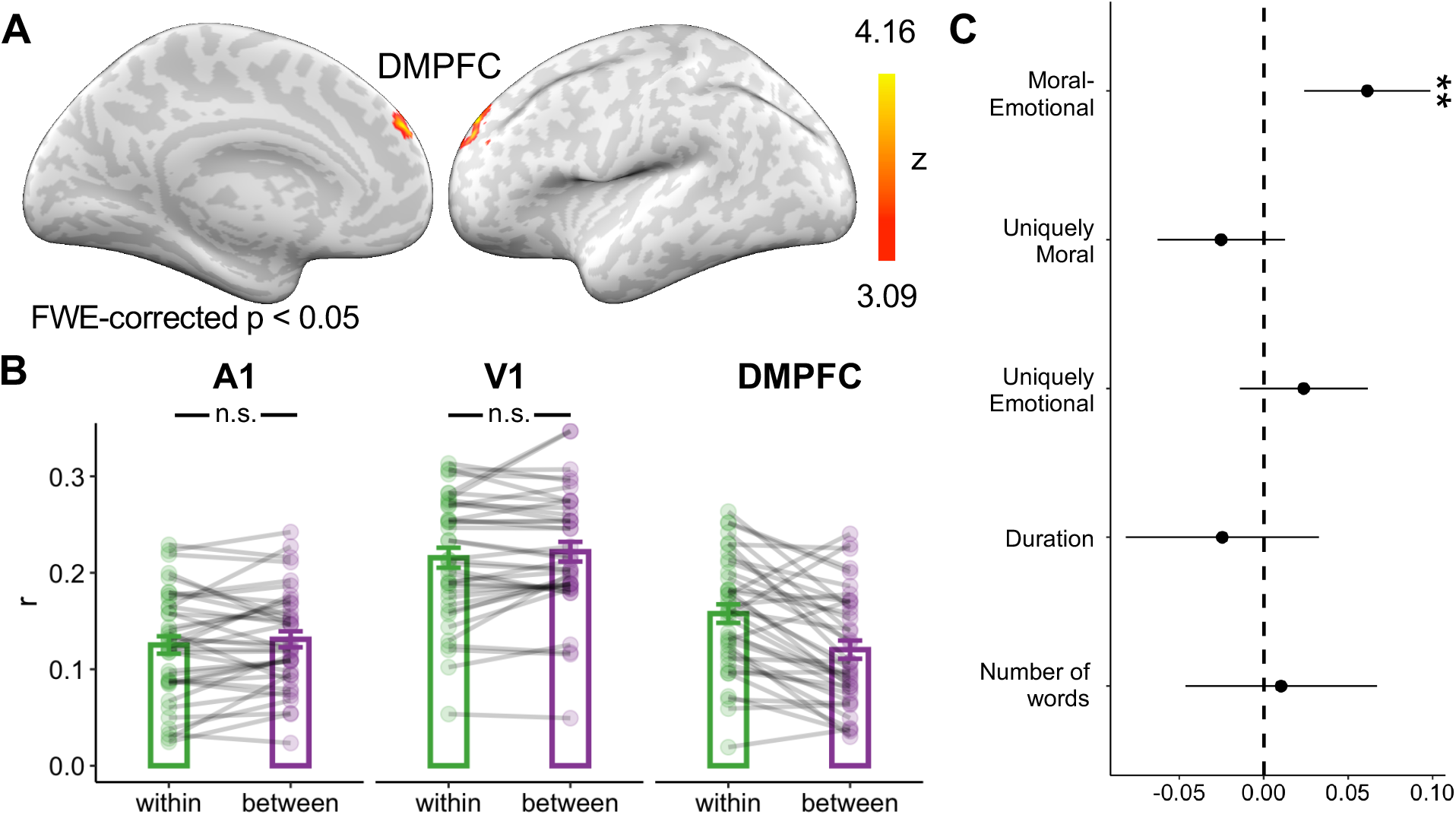
DMPFC timecourse diverges between conservatives and liberals. **A.** Within-group ISC was higher than between-group ISC in the left DMPFC. *p*-values were computed by comparing the observed ISC difference to a null distribution generated using a non-parametric permutation procedure (see Methods). We imposed a FWE cluster-correction threshold of p < 0.05 with a cluster-forming threshold of p < 0.001. Unthresholded map: https://neurovault.org/collections/PKFXOYLX/images/319402/. **B.** Within-group ISC was not significantly different from between-group ISC in both the primary auditory cortex (A1; *t*(37) = - 1.26, *p* = 0.215) and the primary visual cortex (V1; *t*(37) = −1.54, *p* = 0.133). For comparison, we display the ISC values for the DMPFC, but no additional inferences should be drawn based on these plots as the statistical contrast used to identify the DMPFC predetermined a significant difference. Datapoints denote individual participants, error-bars denote between-participant S.E.M. **C.** The use of moral-emotional language was associated with greater neural polarization. Data points indicate regression coefficients with 95% confidence intervals estimated from a linear model. ** < 0.01

### Neural polarization in DMPFC is associated with use of moral-emotional language

To examine the content features that drive neural polarization in the DMPFC, we segmented the twenty-four videos into 86 shorter “segments” (average duration = 24.7s, SD = 5.56s). We averaged DMPFC activity in each segment separately for liberal and conservative-leaning participants to obtain an average liberal timecourse and average conservative timecourse (Fig. S3A). We then computed the absolute difference between the two timecourses as a continuous measure of neural polarization in the DMPFC (Fig. S3B).

We first assessed if moral and emotional content was associated with greater neural polarization in the DMPFC. Using the Moral Emotional dictionary (30), we identified words that relate to both morality and emotions (moral emotional words; e.g., “compassionate”, “violate”), words that relate to morality but not emotions (uniquely moral words; e.g., “ethics”, “principles”) and words that relate to emotions but not morality (uniquely emotional words; e.g., “rewarding”, “fear”).

For each segment, we calculated the percentage of moral-emotional, uniquely moral and uniquely emotional words. We then entered these percentages as predictor variables in the same linear regression model to predict neural polarization in the DMPFC, with the number of words and duration of the segment as additional covariates. Moral-emotional words were associated with greater neural polarization (b = 0.06, 95% CI [0.02, 0.10], *t(*80) = 3.26, *p* = 0.002, Fig. 3C), while uniquely moral (b = −0.02, 95% CI [-0.06, 0.01], *t(*80) = −1.33, *p* = 0.187) and uniquely emotional words (b = 0.02, 95% CI [-0.01,0.06], *t(*80) = 1.24, *p* = 0.218) were not, suggesting that the use of moral-emotional language led to greater polarization in neural responses.

### Data-driven linguistic analysis of content features associated with neural polarization

Next, we assessed the relationship between neural polarization and a broader set of content features. We calculated the percentage of words in each segment that fell into the 47 semantic categories included in the Linguistic Inquiry and Word Count software (LIWC; 31). This allowed us to quantify the extent to which each segment contained words that relate to a variety of semantic content. Using partial least squares (PLS) regression, we examined the relationship between semantic content and neural polarization in the DMPFC. We entered the percentage of words in the 47 LIWC categories and the 3 categories from the Moral Emotional dictionary as predictor variables in a regression model. The duration and number of words in each segment were also included in the model as regressors of no interest.

PLS regression reduces the dimensionality of the predictor variables to a smaller set of latent components that best explains variation in both the predictor and response variables (32). It is commonly used when there are many predictor variables and limited observations, as is the case in our model (i.e. 50 semantic categories predicting neural polarization in 86 segments). We relied on a model that reduced the predictor variables to a single component as it produced a lower cross-validated prediction error than models with additional components (Fig. S4).

Figure 4A shows the regression coefficient for each variable estimated from the model (numerical values reported in Table S2). While no formal test for significance has been developed for the regression coefficients of PLS models, we can estimate a *t*-statistic and *p*-value for each predictor using a jackknife procedure (see Methods) (33). However, as the distribution of the t-statistics for these coefficients is currently unknown, the p-values should be interpreted with caution. Risk-related words (e.g., “threat”, “security”; *t*(85) = 4.10, Bonferroni corrected *p* = 0.005) and moral-emotional words (*t*(85) = 3.45, Bonferroni corrected *p* = 0.044) were the strongest predictors of neural polarization in the DMPFC, suggesting that these two categories of words were most responsible for driving differences between conservative and liberal participants in our experiment. Example segments containing risk-related and moral-emotional words are shown in Fig. 4B and 4C respectively.

**Figure 4.**
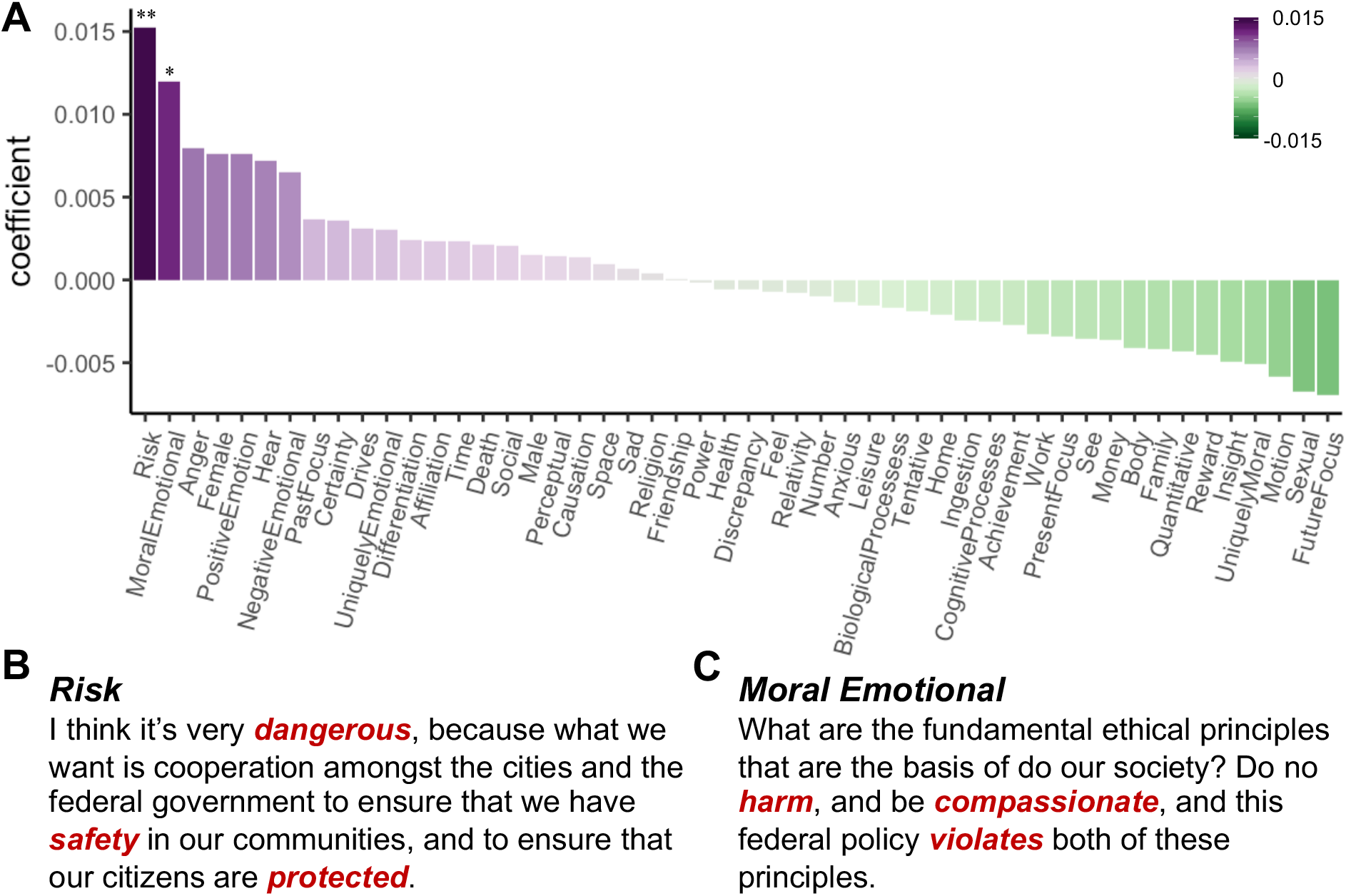
Relationship between semantic content and neural polarization in DMPFC. **A.** Regression coefficients estimated using partial least squares regression. **B.** Red font indicates examples of risk-related words. **C.** Red font indicates examples of moral emotional words. * Bonferroni corrected p < 0.05; ** Bonferroni corrected p < 0.01

### Political differences are associated with divergent frontostriatal connectivity

How might the neural polarization in the DMPFC arise? One possibility is that inputs to the DMPFC are modulated by one’s preexisting political attitudes. That is, even if neural responses in other brain areas were similar between the two groups, differential connectivity to the DMPFC could drive different DMPFC responses to the videos. We ran intersubject functional connectivity (ISFC) analyses to test if connectivity to the DMPFC was modulated by political attitudes.

Functional connectivity (FC) between brain regions is thought to reflect inter-regional communication (34). While conventional FC analyses compute FC as the inter-regional correlation *within* each participant’s brain, ISFC analyses compute the inter-regional correlation *between* brains. In doing so, the ISFC approach filters out within-participant correlations unrelated to stimulus processing (35). ISFC does not imply that there is communication between brains; the technique merely uses a second brain as a model of neural responses to the stimulus from which to compute stimulus-driven correlations between brain regions.

We computed the correlation between each participant’s DMPFC timecourse and a) the timecourse of each voxel averaged over all other participants in the same political group (within-group ISFC) and b) the timecourse of each voxel averaged over all participants in the other political group (between-group ISFC). The correlation between the ventral striatum and the DMPFC was stronger when computed between participants with similar political attitudes than when computed between participants with dissimilar political attitudes (within-group ISFC > between-group-ISFC; Fig. 5), suggesting that covariation between the ventral striatum and the DMPFC was modulated by political attitudes.

**Figure 5.**
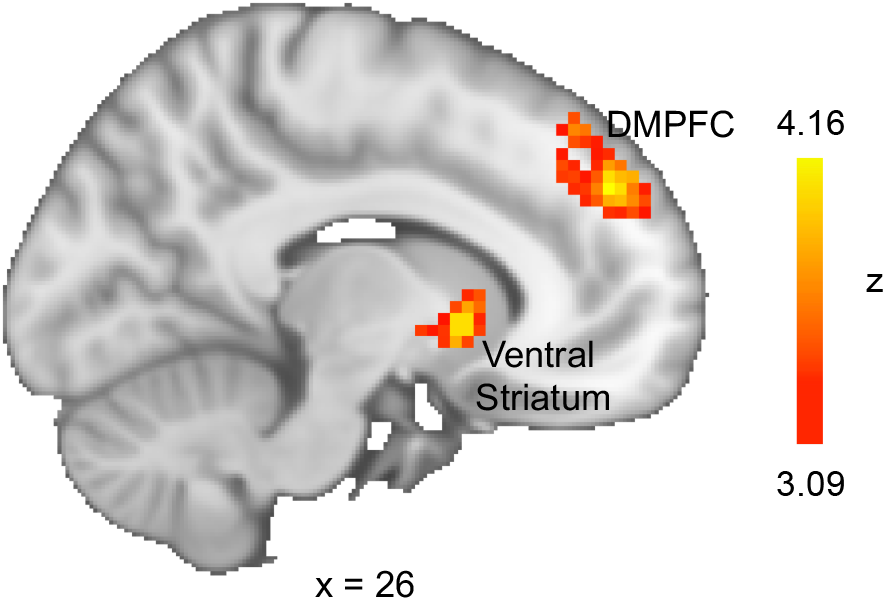
Inter-subject functional connectivity between the ventral striatum and DMPFC was stronger when computed between participants with similar political attitudes. DMPFC was used as a seed region to compute within and between-group ISFC. Statistical map shows voxels where within-group ISFC was higher than between-group ISFC. This analysis also reproduced our earlier result where DMPFC was more correlated within-group than between group. We imposed a FWE cluster-correction threshold of p < 0.05 with a cluster-forming threshold of p < 0.001. Unthresholded map: https://neurovault.org/collections/PKFXOYLX/images/319403/

### Neural similarity to partisan average timecourses predicts video-specific attitude change

After watching each video, participants rated the extent to which the video made them more or less likely to support the relevant policy on a 5-point scale. The responses were recoded such that higher ratings denote attitude change towards the conservative position (e.g., more likely to support the construction of a border wall) while lower ratings denote attitude change towards the liberal position (e.g., less likely to support the construction of a border wall). On average, ratings were higher for conservative participants than liberal participants (M_Conservative_ = 3.16, SE = 0.154, M_Liberal_ = 1.79, SE = 0.105, *t*(31.8) = 7.4, p < 0.001), indicating that participants were more likely to change their attitudes towards the positions held by their respective groups, though there was considerable variability across participants (Fig. S5) and videos (Fig. S6).

We hypothesized that processing a video in a manner that is similar to a particular political group would predict attitude change towards positions held by that group. For each video, we calculated the correlation between each participant’s DMPFC timecourse and (a) the average conservative DMPFC timecourse and (b) the average liberal DMPFC timecourse. The average timecourses were calculated while excluding that participant’s data. We then computed the difference between the two correlations as a measure of whether a participant’s brain activity was more similar to an average conservative (positive values) or an average liberal (negative values).

For a given video, participants whose DMPFC timecourse was more similar to the average conservative or the average liberal participant were more likely to change their attitude towards the conservative position or the liberal position respectively (b = 0.29, SE = 0.12, *t*(876) = 2.43, *p* = 0.015). This analysis was conducted using a linear mixed effects model, and controlled for each participant’s initial attitude towards the policy (see Methods). Similar effects were not observed with participants’ ratings of agreeableness (b = 0.07, SE = 0.136, *t*(867) = 0.51, *p* = 0.614) and credibility (b = −0.07, SE = 0.136, *t*(880) = −0.55, *p* = 0.584) of the videos.

## Discussion

Pre-existing attitudes powerfully influence how individuals respond to political information. In the current work, we combined fMRI and text analysis to study why conservatives and liberals respond differently to the same political content. Activity in the DMPFC diverged between conservative-leaning and liberal-leaning participants watching the same video clips related to immigration policy. This “neural polarization” between the two groups increased with the use of risk-related and moral-emotional language in the videos, highlighting the type of language likely to drive divergent interpretations between the two groups. Neural polarization also tracked subsequent attitude polarization. For each video, participants with DMPFC activity timecourses more similar to that of conservative-leaning participants became more likely to support the conservative position. Conversely, those with DMPFC activity timecourses more similar to that of liberal-leaning participants became more likely to support the liberal position. These results suggest that divergent interpretations of the same information are associated with increased attitude polarization. Together, our findings describe a neural basis for partisan biases in processing political information and their effects on attitude change.

Our approach builds on earlier findings showing that the neural similarity between individuals watching or listening to a stimulus reflects similarity in how the stimulus is processed by a particular brain region (18, 36, 37). In our dataset, neural responses in the primary sensory cortices were not significantly different between conservative-leaning and liberal-leaning participants, suggesting that political attitudes did not alter sensory processing. Neural responses between the two groups differed only in the DMPFC, a brain region that has been associated with the interpretation of narrative stimuli (20, 21, 29). For example, a previous study found that neural responses in the DMPFC diverged between participants manipulated to have different interpretations prior to listening to an ambiguous story (21). Here, we did not manipulate participants to have different interpretations. Instead, participants’ political attitudes served as intrinsic “priors” that biased how they interpreted the content of the videos.

The DMPFC has been implicated in a broad range of complex cognitive functions, including episodic memory retrieval, impression formation and reasoning about other people’s mental states (38–40). One account that integrates over these disparate findings is that the DMPFC is involved in the construction of situation models - mental representations of the actors, actions and objects in an event, as well as their spatial, temporal and causal relationships (41–43). The divergence in DMPFC activity between conservative-leaning and liberal-leaning participants might thus reflect the two groups constructing different situation models of the events depicted in the videos. To better understand how conservative-leaning and liberal-leaning participants interpreted the videos differently, we analyzed the content of the videos to find semantic features that would be associated with greater neural polarization in the DMPFC.

Risk-related and moral-emotional words were most likely to drive neural polarization in the DMPFC. These results are consistent with two major lines of work in political psychology. First, several prominent theories have proposed that conservatives and liberals exhibit different levels of threat sensitivity (44–46). Risk-related words are often used to describe potential threats, which would trigger different responses in the two groups. Second, conservatives and liberals are thought to adopt different moral frameworks, and thus see the world through distinct “moral lenses” (22–24). As such, what conservatives consider a moral transgression may seem perfectly acceptable to liberals, and vice versa. Taken together, these theories would predict that conservatives and liberals would have different interpretations of what is a threat and what is morally praiseworthy or blameworthy. This would explain why the neural polarization in the DMPFC was greater in parts of the videos with more risk-related and moral-emotional words.

Participants’ interpretation of the videos would likely modulate if and how the videos changed their attitudes. To examine the relationship between video interpretation and attitude change, we used DMPFC activity as a neural model of how participants interpreted each video. Specifically, we averaged the DMPFC timecourse separately for conservative-leaning and liberal-leaning participants. These average timecourses reflect how the typical conservative-leaning or liberal-leaning participants processed each video. We thus computed the similarity between each participant’s DMPFC timecourse and the two average timecourses to obtain a neural metric of whether a participant’s interpretation was more similar to the conservative interpretation or the liberal interpretation. For a given video, neural similarity to a particular group was associated with attitude change towards the positions held by that group. This finding suggests that adopting the liberal interpretation of a video biased participants towards the liberal position while adopting the conservative interpretation biased participants towards the conservative position.

How might the differential response in the DMPFC arise in the brain? One possibility is that inputs to the DMPFC were modulated by political attitudes. Consistent with this hypothesis, we found that inter-subject functional connectivity between the ventral striatum and the DMPFC was stronger between individuals with similar political attitudes. The ventral striatum is commonly associated with the processing of affective valence (i.e. whether an experience is positive or negative) (47, 48). Our results suggest that the propagation of valence information from the ventral striatum to the DMPFC was modulated by one’s political attitudes. This biases the DMPFC response, giving rise to the divergence in DMPFC activity between participants with dissimilar political attitudes. The temporal resolution of fMRI data, however, does not allow us to make claims about the directionality of influence. As such, this interpretation is speculative and future studies will be needed to clarify the role of frontostriatal connectivity in modulating DMPFC responses.

It is important to test if our results would generalize to other polarizing issues, such as abortion, gun control and climate change. Given that political differences on these issues are also associated with differences in threat-perception and moral values (49, 50), we predict that this would be the case. Future studies should also examine the extent to which our results would apply to polarized groups outside the American political context. Recent work found that political messages with more moral-emotional words were more likely to spread on online social networks (30, 51). This spread, however, was contained within networks of individuals who share similar political attitudes and can contribute to further attitude polarization. The polarized neural responses observed in our experiment suggests a neural precursor to the biased diffusion of political information. In particular, messages that induce greater polarization between groups might also be messages that are more likely to spread in a polarized manner. This hypothesis can be tested by measuring the neural responses of participants reading messages, and analyzing whether, and with whom, the messages are subsequently shared.

Our work introduces a novel multimethod approach to study the political brain under naturalistic conditions. Using this approach, we identified a neural correlate of the biased processing of political information, as well as the content features most likely to be processed in a biased manner. Divergent interpretations, as indexed by neural activity, were in turn associated with attitude change in response to the videos. Future work can combine neuroimaging data with machine learning methods in natural language processing to build semantic models of how political content is interpreted, and inform interventions aimed at aligning interpretations between conservative and liberals.

## Methods

### Participants

Forty participants were recruited from the Stanford community using an online human participant management platform (SONA systems). Participants interested in the study first indicated their age and sex, as well as their support for the six immigration policies (see *Experimental Task*) on an online questionnaire. We made an effort to recruit participants with varying immigration attitudes. As the pool of participants leaned liberal, this required oversampling of participants with conservative-leaning attitudes. Participants received $50 for participating in the two-hour experiment. Data from two participants were discarded because of excessive head motion (> 3mm) during one or more scanning sessions, yielding an effective sample size of thirty-eight participants (23 male, 15 female, ages 19-57, mean age = 31.3). All participants provided written, informed consent prior to the start of the study. Experimental procedures were approved by the Stanford Institutional Review Board.

### Experimental Task

Participants were scanned using fMRI as they watched twenty-four videos (total duration = 35min 26s, divided into four runs) on six immigration policies: i. *Border Wall*: the construction of a wall along the U.S.-Mexico border to reduce illegal immigration, ii. *Work Authorization*: allowing illegal/undocumented immigrants to work legally in the U.S. without fear of deportation, iii. *Refugee Ban*: banning refugees from majority-Muslim countries from entering the U.S., iv. *Healthcare Provision*: allowing the use of federal funds to pay for emergency healthcare for undocumented/illegal immigrants, v. *Dream Act*: providing a pathway to citizenship for undocumented individuals brought into the U.S. illegally as children, vi. *Sanctuary cities:* cutting federal funding to sanctuary cities unless the cities agree to fully cooperate with the U.S. immigration and customs enforcement (ICE). All videos were obtained from youtube.com, and were selected to represent both liberal and conservative viewpoints. We provide the duration and a one-sentence description of each video in Table S3.

Participants were instructed to watch the videos as they would watch television at home. At the end of each video, participants were asked to rate on a five-point scale how much they agreed with the general message of the video (*agreeableness*), how credible was the information presented in the video (*credibility*), and the extent to which the video made them more or less likely to support the policy in question (*change*).

### Pre- and post-experimental measures

Prior to being scanned, participants first completed a questionnaire on which they indicated their support for the six immigration policies (7-point scale from Strongly Not Support to Strongly Support), their political orientation (7-point scale from Extremely Liberal to Extremely Conservative) and political affiliation (Strong Democrat, Moderate Democrat, Independent, Moderate Republican, Strong Republican). At the end of the scanning session, participants completed the same questionnaire, and also provided information about their annual household income (in $10,000 increments from 0 to $100000, $100000 to $150000, More than $1500000), and education levels (a. Less than high school, b. High School/GED, c. Some college, d. 2-year college degree (Associates), e. 4-year college degree (BA, BS), f. Master’s Degree, g. Doctorate or Professional Degree).

### fMRI data acquisition and preprocessing

MRI data were collected using a 3T General Electric MRI scanner. Functional images were acquired in interleaved order using a T2*-weighted echo planar imaging (EPI) pulse sequence (46 transverse slices, TR=2s, TE=25ms, flip angle=77°, voxel size 2.9 mm^3^). Anatomical images were acquired at the start of the session with a T1-weighted pulse sequence (TR = 7.2ms, TE = 2.8ms, flip angle=12°, voxel size 1 mm^3^). Image volumes were preprocessed using FSL/FEAT v.5.98 (FMRIB software library, FMRIB, Oxford, UK). Preprocessing included motion correction, slice-timing correction, removal of low-frequency drifts using a temporal high-pass filter (100ms cutoff), and spatial smoothing (4-mm FWHM). Functional volumes were first registered to participants’ anatomical image (rigid-body transformation with 6 degrees of freedom) and then to a template brain in Montreal Neurological Institute (MNI) space (affine transformation with 12 degrees of freedom).

### Intersubject correlation analyses

We reordered and concatenated the neural data such that the twenty-four videos were ordered in the same sequence for all participants. We then computed the *one-to-average* intersubject correlation across the entire sample. For each participant, we computed the Pearson correlation between the activity timecourse of a voxel with the activity timecourse at the same voxel averaged across all other participants. This procedure was repeated for all voxels and averaged over participants to obtain a map of average *r*-values.

We assessed statistical significance using a non-parametric permutation test. For each voxel, we computed the *t*-statistic testing if the average *r*-value was greater than zero. To generate a null-distribution, we flipped the sign of the *r*-values for a random subset of participants and recomputed the *t*-statistic. This procedure was repeated 100,00 times. The *p-value* was computed as the proportion of the null distribution that was more positive than the observed *t*-statistic. We thresholded the statistical map for voxels that survive correction for multiple comparisons to control for false discovery rate (FDR) using the two-stage Benjamini, Krieger, & Yekutieli procedure (52) (q < 0.001).

### Within-group vs. between-group analyses

For each participant, we calculated an immigration attitude score by tallying their support for the six immigration policies. The responses were coded such that higher ratings correspond to a stronger conservative-leaning. We then performed a median split to categorize participants into conservative and liberal participants. We ran two-sample t-tests to test for group differences age, head-motion (framewise displacement), education and household income, and a chi-square test to test for group differences in sex. We also computed the Spearman correlation between the continuous immigration attitude scores and age, head-motion, education and household income.

We searched for voxels where the timecourse of activity was more similar within-group than between-group. For each participant, we computed the within-group ISC as the voxel-wise correlation with the average timecourse of all other participants in the group (i.e. correlation between the activity of a liberal participant and the average activity of all other liberal participants; correlation between the activity of a conservative participant and the average activity of all other conservative participants). Conversely, we computed the between-group ISC as the voxel-wise ISC with the average timecourse of participants in the other group (i.e. correlation between the activity of a liberal participant and the average activity of conservative participants; correlation between the activity of a conservative participant, and the average activity of liberal participants). For each participant and each voxel, we computed the difference between the within-group ISC and between-group ISC. This difference was then averaged across all participants to obtain a map of average difference in *r*-values, which reflects the extent to which the activity timecourse was more similar within groups than between groups.

We used a non-parametric permutation test to assess statistical significance of the difference map. For each voxel, we computed the *t*-statistic testing if the average difference in *r* was greater than zero. To generate a null-distribution, we flipped the sign of the difference in *r* for a random subset of participants and recomputed the *t*-statistic. This procedure was repeated 100,00 times. The *p-value* was computed as the proportion of the null distribution that was more positive than the observed *t-*statistic. We imposed a family-wise error cluster-correction threshold of p < 0.05 using Gaussian Random Field theory, with a cluster-forming threshold of p < 0.001.

### Intersubject functional connectivity analyses

A DMPFC region of interest (ROI) was defined as the voxels that survived correction in the within-group ISC > between-group ISC contrast. For each participant, we extracted the average timecourse in this ROI. For each voxel, we then computed the correlation between the average DMPFC timecourse and the voxel-wise timecourse averaged over all other participants in the same political group (within-group ISFC), and that averaged over all participants in the other political group (between-group ISFC). For each participant and each voxel, we computed the difference between the within-group ISFC and between-group ISFC. This difference was then averaged across all participants to obtain a map of average difference in *r*, which reflects the extent to which the activity timecourse was more correlated with average DMPFC activity of participants with similar immigration attitudes and that of participants with opposite immigration attitudes.

We used a non-parametric permutation test to assess statistical significance of the difference map. For each voxel, we computed the *t*-statistic testing if the average difference in *r* was greater than zero. To generate a null-distribution, we flipped the sign of the difference in *r* for a random subset of participants and recomputed the *t*-statistic. This procedure was repeated 100,00 times. The *p-value* was computed as the proportion of the null distribution that was more positive than the observed *t-*statistic. We imposed a family-wise error cluster-correction threshold of p < 0.05 using Gaussian Random Field theory, with a cluster-forming threshold of p < 0.001.

### Linguistic content analyses (linear regression)

To examine how the content of the videos relate to the differences in neural processing between liberals and conservatives, we transcribed the audio for all videos and segmented the videos into smaller segments corresponding to pauses in speech or a switch between different speakers (84 segments in total, range = 12-38s, average duration = 24.7s, SD = 5.56s). For each segment, we used the Linguistic Inquiry and Word Count software (LIWC) (31) to count the proportion of words that fall into three distinct word categories: Moral Emotional words, Uniquely Moral words, Uniquely Emotional words. *Moral Emotional* words are words that appear in both the Moral Foundations dictionary (24) and in the Affect dictionary included with the LIWC software. *Uniquely Moral* words are words that appear in the Moral Foundations dictionary but not the Affect dictionary, while *Uniquely Emotional* words are words that appear in the Affect dictionary but not the Moral Foundations dictionary. Previous work has shown that this procedure generates three categories of words with high discriminant validity - moral-emotional words are rated as more moral than uniquely emotional words, and more emotional than uniquely moral words (30).

We defined a DMPFC region of interest (ROI) as the cluster of voxels where activity was more similar within groups than between groups. For each segment, we computed the average DMPFC activity separately for liberal and conservative participants. We then obtained the absolute difference between the average activity of the two groups as a measure of neural polarization. Using a linear model, we tested if the proportion of moral emotional words, uniquely moral words, and uniquely emotional words in a segment would predict the magnitude of neural polarization, controlling for the number of words and the duration of the segment. All variables were mean-centered and scaled by two standard deviation prior to being entered into the model to facilitate the comparison of the resulting regression coefficients on a common scale (53).

### Linguistic content analyses (partial least squares regression)

We then used partial least squares regression to examine the relationship between neural polarization in the DMPFC and a broader set of semantic categories. For each segment, we calculated the proportion of words in the 47 semantic categories included as part of the LIWC software and the 3 categories from the Moral Emotional dictionary. We entered the proportion of words in the 50 semantic categories as predictor variables in a partial least squares regression model to predict neural polarization. The model was fitted using the *pls* package in R (32). The duration and number of words in each segment were also included in the model as regressors of no interest.

We used a leave-one-segment-out-cross-validation procedure to select the optimal number of components for the model. Briefly, the model is fit to all but one segment and then tested on the left-out segment. This procedure is repeated 84 times, leaving out a different segment each time. Model performance was computed as the root mean squared error of prediction (RMSEP) of the left-out segments. The cross-validation procedure was performed separately for models where the predictors were reduced to a different number of components (1 to 5). The regression coefficients for each predictor was then computed using the model with the lowest RMSEP.

A t-statistic for each predictor was estimated using a jackknife procedure, and a *p*-value computed with *n* - 1 degrees of freedom where *n* is the number of segments (33). The *p*-value was then Bonferroni corrected for 50 tests (corrected *p*-value = *p-*value x number of tests). We note, however, that the distribution of the *t-*statistics from a partial least squares regression is currently unknown, and as such, these *p*-values should be interpreted with caution.

### Predicting video-specific attitude change

For each video, we computed the correlation between each participant’s DMPFC timecourse and the average conservative and average liberal DMPFC timecourses. The average timecourses were computed while excluding that participant’s data. We then computed the difference between the two correlations as a measure of whether a participants’ brain activity was more similar to an average conservative or an average liberal. Positive values indicate greater similarity to the average conservative timecourse while negative values indicate greater similarity to the average liberal timecourse.

We ran three separate linear mixed effects model to predict video-specific ratings *agreeableness*, *support* and *change* (see *Experimental Task*) from the neural similarity to conservative vs. liberal participants. Participants initial attitude towards the policy mentioned in the video was included as a regressor of no interest. The models were estimated using the lmer function in the lme4 package in R (54), with *p*-values computed from *t*-tests with Satterthwaite approximation for the degrees of freedom as implemented in the lmerTest package (55).

## Data availability

Data will be uploaded and made publicly available on https://openneuro.org/ upon publication.

## Author contributions

Y.C.L. collected and analyzed the data. All authors contributed to the design of the study and writing of the manuscript.

## Acknowledgments

We thank Courtney Gao, Yiyu Wang and Gloria Wong for assistance with stimuli creation, video transcription and data collection, and all members of the Stanford Social Neuroscience Lab for helpful discussion. This research was supported by the Army Research Office (grant number: W911NF1610172 to J.Z.).

