## Supplemental Information for "Conservative and liberal attitudes drive polarized neural responses to political content"

### Supplementary Results

**Online pre-test.** 300 US-based participants (179 male, 120 female, 1 other; 21-72 years of age, mean age = 35.03 years) were recruited on the Amazon Mechanical Turk (AMT) online platform. Participants were first asked to indicate their political orientation on a 1-7 scale ($orientation_{score}$: 1 = extremely liberal, 4 = moderate, 7 = extremely conservative), and their support for each of the six immigration policies (1 = strongly not support, 7 = strongly support). 179 participants identified as liberal ($orientation_{score} < 4$), 49 participants identified as moderate ($orientation_{score} = 4$) and 72 participants as conservative ($orientation_{score} > 4$).

We tested if support for each policy differed between conservatives and liberals. Statistical significance was assessed using a Welch Two Sample t-test, and results are depicted graphically in Fig. S1. Relative to conservative participants, liberal participants were more likely to support allowing illegal/undocumented immigrants to work legally in the US ($M_{liberal} = 4.83$, $SE_{liberal} = 0.13$, $M_{conservative} = 2.12$, $SE_{conservative} = 0.19$, $t(142.1) = 11.5$, $p < 0.001$), allowing the use of federal funds to pay for emergency healthcare for undocumented/illegal immigrants ($M_{liberal} = 4.69$, $SE_{liberal} = 0.14$, $M_{conservative} = 2.58$, $SE_{conservative} = 0.23$, $t(121.6) = 7.81$, $p < 0.001$), and providing a pathway to citizenship for undocumented individuals brought into the U.S. illegally as children ($M_{liberal} = 6.08$, $SE_{liberal} = 0.10$, $M_{conservative} = 4.01$, $SE_{conservative} = 0.23$, $t(97.9) = 8.01$, $p < 0.001$).

Relative to liberal participants, conservative participants were more likely to support funding the construction of a wall along the U.S.-Mexico border to reduce illegal immigration ($M_{conservative} = 5.04$, $SE_{conservative} = 0.24$, $M_{liberal} = 1.73$, $SE_{liberal} = 0.10$, $t(98.7) = 12.7$, $p < 0.001$), banning refugees from Muslim-majority countries from entering the country ($M_{conservative} = 4.66$, $SE_{conservative} = 0.25$, $M_{liberal} = 2.11$, $SE_{liberal} = 0.11$, $t(104.2) = 9.33$, $p < 0.001$), cutting federal funding to sanctuary cities unless the cities agree to fully cooperate with the U.S. immigration and customs enforcement ($M_{conservative} = 5.29$, $SE_{conservative} = 0.25$, $M_{liberal} = 2.59$, $SE_{liberal} = 0.14$, $t(135.6) = 10.39$, $p < 0.001$).

**Supplementary Figures**

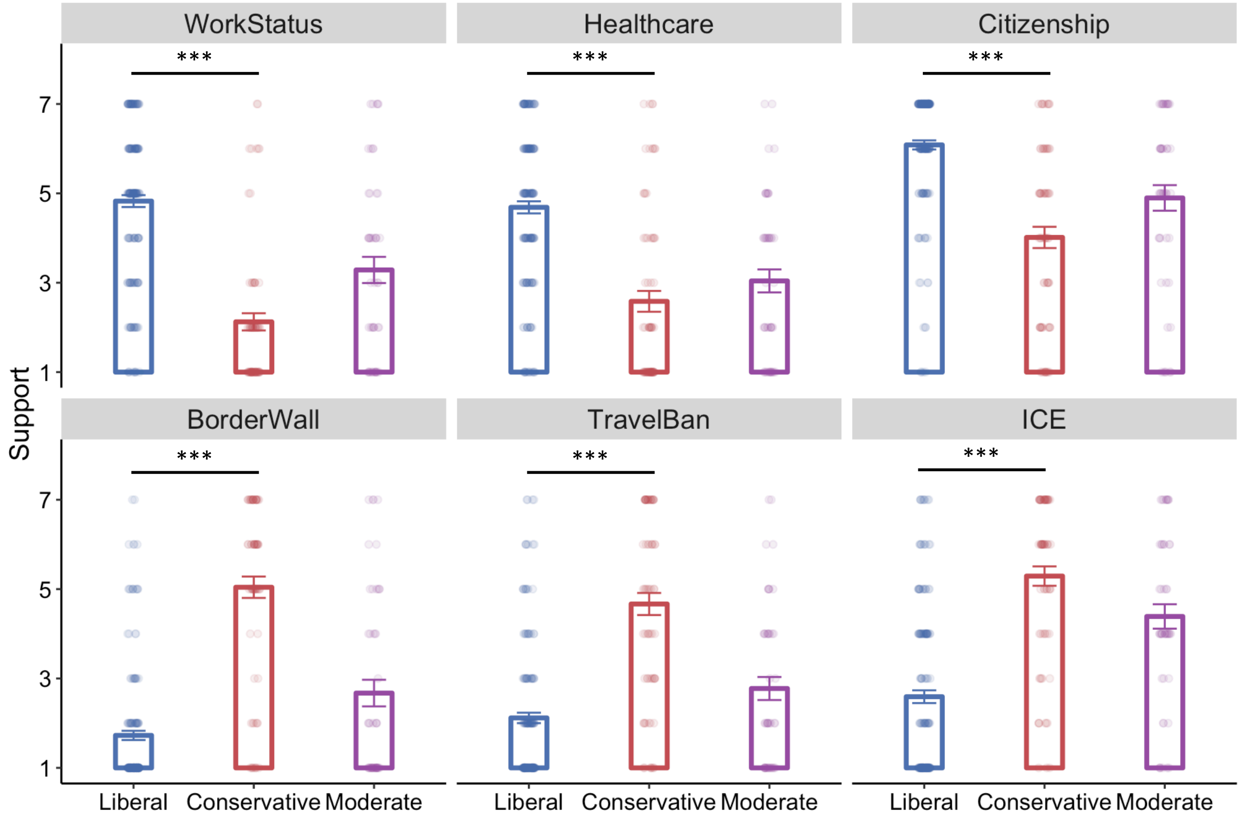


**Figure S1.** Liberals and conservatives differ significantly on their support for each of the six policies. See Fig. 1B and Supplemental Results for full description of each policy. Data points indicate individual participants' support for the policy with horizontal jitter added for clearer visualization. Support by participants identifying as moderate are shown in purple for comparison. Error bars indicate SEM. *** p < 0.001

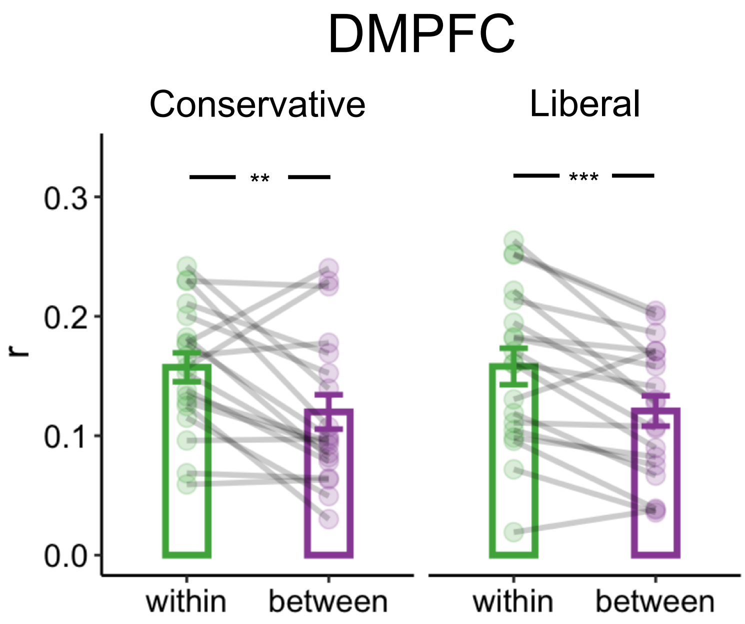


**Figure S2.** Within-group ISC was higher than between-group ISC in both conservative (*t(18)* = 3.01, p = 0.007) and liberal participants (*t(18)* = 4.57, *p* < 0.001). Furthermore, political orientation did not moderate the within vs. between group ISC difference (*t(36)* = 0.007, *p* = 0.994) . ** p < 0.01, *** p < 0.001.

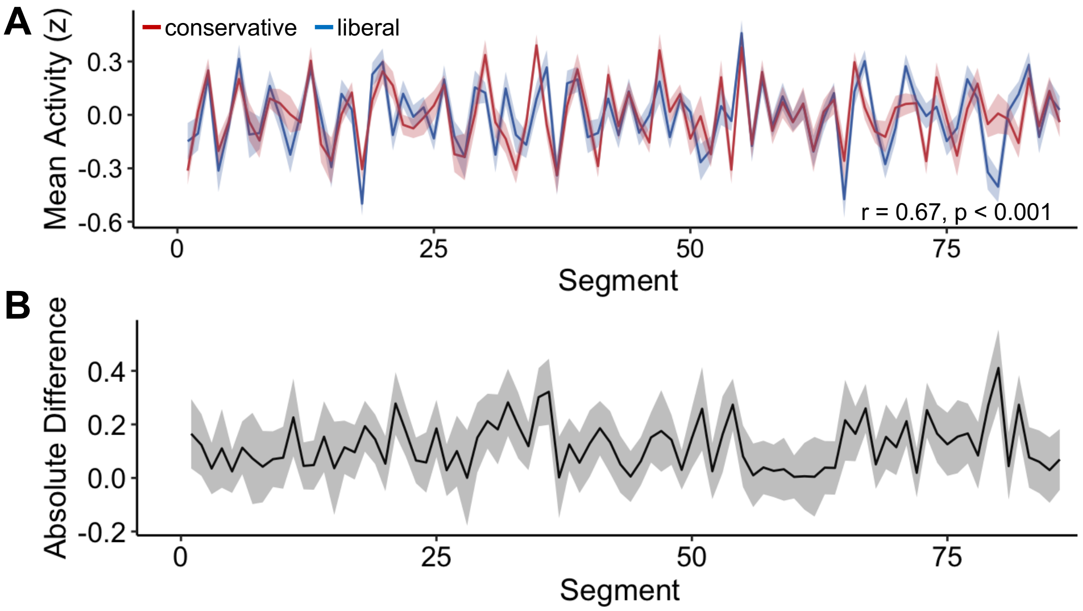


**Figure S3.** Neural polarization in the DMPFC. **A.** Average DMPFC timecourse of conservative (red) and liberal (blue) participants over the 86 segments. The two timecourses were moderately correlated (r = 0.66, $p$ < 0.001). **B.** Absolute difference between average conservative and average liberal DMPFC activity. Shaded errors indicate SEM.

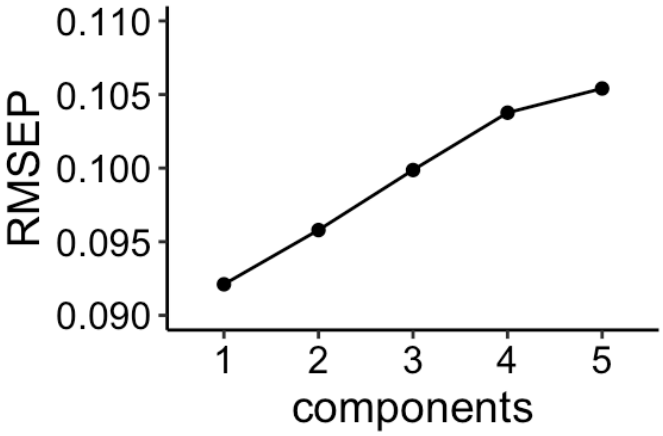


**Figure S4.** Root mean square error of prediction (RMSEP) calculated from leave-one-segment-out cross-validation was lowest for a model with 1 component.

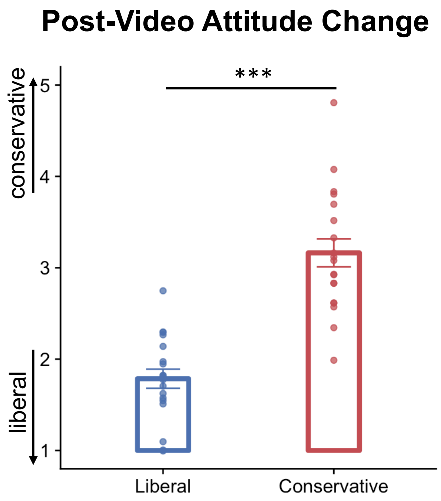


**Figure S5. Ratings post-video attitude changes averaged across videos separately for liberal-leaning (blue) and conservative-leaning (red) participants.** Higher ratings denote attitude change towards the conservative position while lower ratings denote attitude change towards the liberal position. Datapoints indicate average rating for individual participants. *** $p < 0.001$

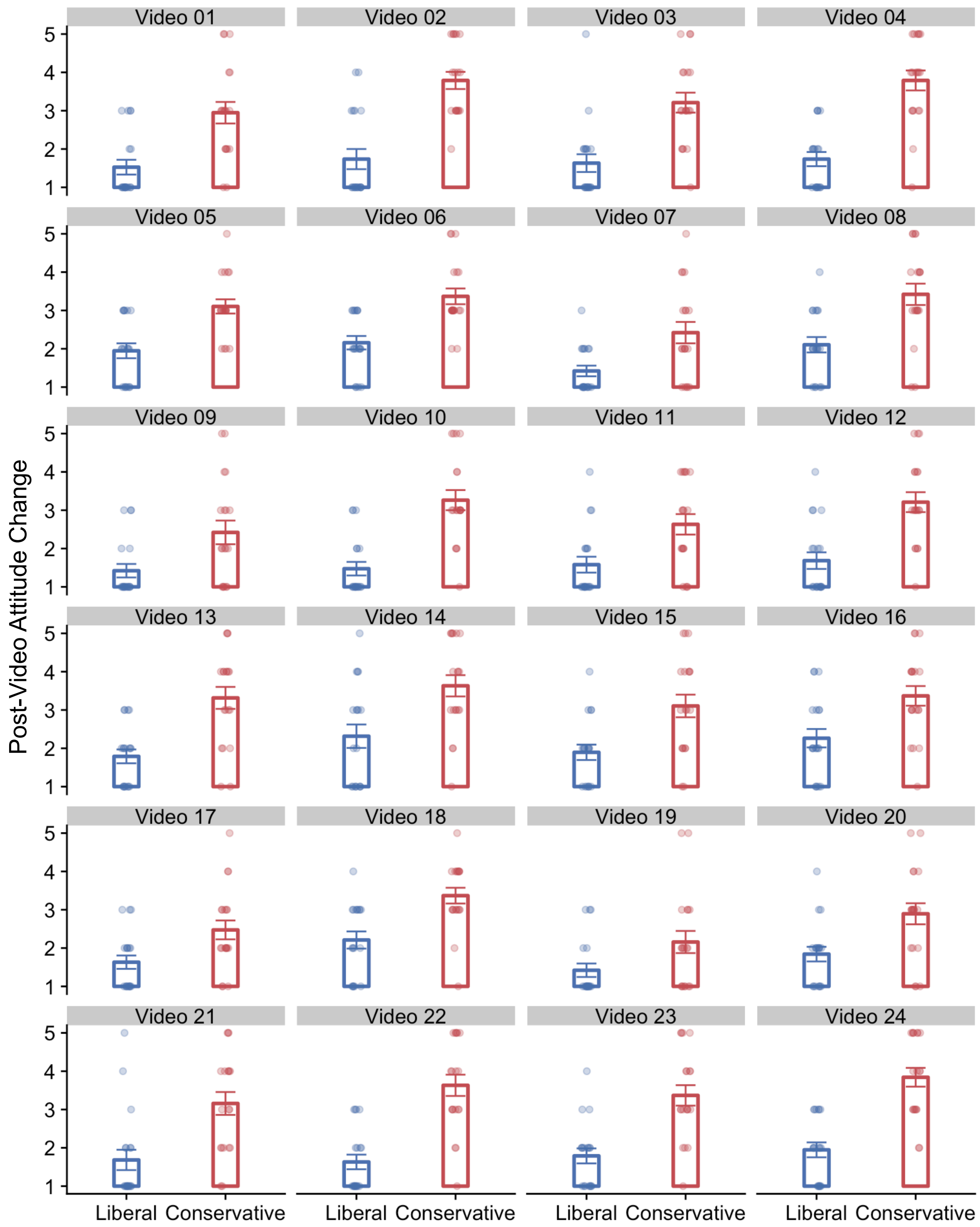


**Figure S6. Post-video attitude change for each video separately for liberal-leaning (blue) and conservative-leaning (red) participants.** Higher ratings denote attitude change towards the conservative position while lower ratings denote attitude change towards the liberal position. Datapoints indicate rating for each participant with horizontal jitter added for clearer visualization.

### Supplementary Tables

**Table S1. Immigration attitude was not related to potentially confounding variables.** Liberal-leaning and conservative-leaning participants did not differ significantly on age, sex, income, education and head-motion in the scanner (FD: framewise displacement). A paired t-test was used to test for differences between groups for all variables except sex, where a chi-square test was used instead. We also computed the Spearman correlation coefficient to test for continuous relationships between immigration attitude and each variable.[1]chi-square statistic.

| | | | Categorical | | Continuous | |
|---|---|---|---|---|---|---|
| | **Liberal** | **Conservative** | ***t*** | ***p*** | **Spearman *rho*** | ***p*** |
| Mean Age | 30.1 (2.1) | 33.0 (3.1) | -0.78 | 0.44 | 0.16 | 0.34 |
| Sex | 9F 10M | 8F 11M | 0.11[1] | 0.74 | - | - |
| Median Income | $80,000-$89,000 | $80,000-$89,000 | -0.38 | 0.70 | 0.09 | 0.56 |
| Median Education | 4-year college degree | 4-year college degree | 1.02 | 0.31 | -0.01 | 0.97 |
| Head Motion: Mean FD (SE) | 0.34 (0.19) | 0.27 (0.11) | 1.46 | 0.16 | -0.01 | 0.95 |

**Table S2. Regression coefficients from partial least squares regression predicting neural polarization in the DMPFC from semantic categories.** t-statistics and p-values were computed using a jackknife procedure, and Bonferroni corrected for 50 tests.

| Regressor | Coefficient | SE | *t*(85) | *p* | Corrected *p* |
|---|---|---|---|---|---|
| Risk | 0.015 | 0.004 | 4.099 | 0.00009 | 0.005 |
| Moral Emotional | 0.012 | 0.003 | 3.449 | 0.001 | 0.044 |
| Anger | 0.008 | 0.004 | 1.926 | 0.057 | 1 |
| Female | 0.008 | 0.004 | 2.047 | 0.044 | 1 |
| Positive Emotion | 0.008 | 0.004 | 1.904 | 0.06 | 1 |
| Hear | 0.007 | 0.004 | 1.734 | 0.086 | 1 |
| Negative Emotion | 0.006 | 0.004 | 1.785 | 0.078 | 1 |
| Past Focus | 0.004 | 0.004 | 0.859 | 0.393 | 1 |
| Certainty | 0.004 | 0.005 | 0.773 | 0.441 | 1 |
| Drives | 0.003 | 0.003 | 0.9 | 0.371 | 1 |
| Uniquely Emotional | 0.003 | 0.004 | 0.767 | 0.445 | 1 |
| Differentiation | 0.002 | 0.004 | 0.54 | 0.59 | 1 |
| Affiliation | 0.002 | 0.005 | 0.454 | 0.651 | 1 |
| Time | 0.002 | 0.004 | 0.617 | 0.539 | 1 |
| Death | 0.002 | 0.004 | 0.536 | 0.594 | 1 |
| Social | 0.002 | 0.005 | 0.446 | 0.657 | 1 |
| Male | 0.002 | 0.005 | 0.331 | 0.742 | 1 |
| Perceptual | 0.001 | 0.004 | 0.333 | 0.74 | 1 |
| Causation | 0.001 | 0.004 | 0.314 | 0.755 | 1 |
| Space | 0.001 | 0.004 | 0.255 | 0.8 | 1 |
| Sad | 0.001 | 0.002 | 0.306 | 0.76 | 1 |
| Religion | 0 | 0.003 | 0.131 | 0.896 | 1 |
| Friendship | 0 | 0.004 | 0.019 | 0.985 | 1 |
| Power | 0 | 0.004 | -0.049 | 0.961 | 1 |
| Health | -0.001 | 0.004 | -0.134 | 0.894 | 1 |
| Discrepancy | -0.001 | 0.004 | -0.138 | 0.89 | 1 |
| Feel | -0.001 | 0.004 | -0.171 | 0.865 | 1 |
| Relativity | -0.001 | 0.004 | -0.197 | 0.844 | 1 |
| Number | -0.001 | 0.003 | -0.341 | 0.734 | 1 |
| Anxious | -0.001 | 0.005 | -0.281 | 0.779 | 1 |
| Leisure | -0.002 | 0.004 | -0.409 | 0.684 | 1 |
| Biological Processes | -0.002 | 0.004 | -0.423 | 0.673 | 1 |
| Tentative | -0.002 | 0.003 | -0.613 | 0.542 | 1 |
| Home | -0.002 | 0.005 | -0.434 | 0.665 | 1 |
| Ingestion | -0.002 | 0.003 | -0.905 | 0.368 | 1 |
| Cognitive Processes | -0.003 | 0.004 | -0.697 | 0.488 | 1 |
| Achievement | -0.003 | 0.003 | -0.9 | 0.37 | 1 |
| Work | -0.003 | 0.003 | -0.969 | 0.335 | 1 |
| Present Focus | -0.003 | 0.004 | -0.861 | 0.392 | 1 |
| See | -0.004 | 0.005 | -0.785 | 0.434 | 1 |
| Money | -0.004 | 0.003 | -1.062 | 0.291 | 1 |
| Body | -0.004 | 0.004 | -1.127 | 0.263 | 1 |
| Family | -0.004 | 0.004 | -1.054 | 0.295 | 1 |
| Quantitative | -0.004 | 0.003 | -1.425 | 0.158 | 1 |
| Reward | -0.005 | 0.004 | -1.196 | 0.235 | 1 |
| Insight | -0.005 | 0.004 | -1.303 | 0.196 | 1 |
| Uniquely Moral | -0.005 | 0.004 | -1.25 | 0.215 | 1 |
| Motion | -0.006 | 0.004 | -1.443 | 0.153 | 1 |
| Sexual | -0.007 | 0.003 | -1.963 | 0.053 | 1 |
| Future Focus | -0.007 | 0.003 | -2.098 | 0.039 | 1 |

**Table S3. Duration and description of videos included in the study**

| Video No. | Issue | Duration | Description |
|---|---|---|---|
| 1 | Border Wall | 1:10 | CNN interview with Senator Dick Durbin (D) on federal funding for a wall on the US-Mexico border |
| 2 | Border Wall | 0:50 | BBC news showing an existing barrier between California and Mexico, followed by an interview with a Border Patrol Agent who supports the construction of a wall along the US-Mexico border |
| 3 | Border Wall | 1:14 | Animated public service announcement on why a wall along the US-Mexico border is unlikely to reduce illegal immigration |
| 4 | Border Wall | 1:47 | Fox Business interview with a researcher from the Center for Immigration Studies on why a wall along the US-Mexico border would reduce illegal immigration |
| 5 | Work Authorization | 1:00 | UC Davis Economics professor explaining how immigrants push Americans towards better paying jobs |
| 6 | Work Authorization | 1:04 | CBS News segment with panelist explaining how immigrants suppress the wages of American workers |
| 7 | Work Authorization | 1:30 | Video clip from President Obama's November 2014 remarks on why he will sign an executive order allowing work authorization of undocumented immigrants |
| 8 | Work Authorization | 1:43 | Animated public service announcement on how illegal immigration results in unemployment and underemployment of American workers |
| 9 | Refugee Ban | 1:55 | Vox video clip with information on the Syrian refugee crisis, followed by criticism of President Trump's executive order to temporarily suspend immigrants and refugees from 7 Muslim-majority countries |
| 10 | Refugee Ban | 1:26 | Local news segment (Buffalo, NY) showing clips from protests against President Trump's executive order to temporarily suspend immigrants and refugees from 7 Muslim-majority countries, followed by clips of individuals voicing their support for the order. |
| 11 | Refugee Ban | 1:48 | CNN journalist Fareed Zakaria providing statistics suggesting that there is "no rational basis" for President Trump's executive order to temporarily suspend immigrants and refugees from 7 Muslim-majority countries |
| 12 | Refugee Ban | 1:57 | Local news segment (Oakland, CA) showing a gathering of Americans who support President Trump's executive order to temporarily suspend immigrants and refugees from 7 Muslim-majority countries, followed by clips of protests against the order at airports. |
| 13 | Healthcare Provision | 1:34 | Animated public service announcement describing how undocumented immigrants pay substantial taxes and should be entitled to healthcare benefits |
| 14 | Healthcare Provision | 1:31 | Fox Business segment with former Arizona governor Jan Brewer criticizing California's petition to expand the Affordable Care Act to include undocumented immigrants, followed by clips showing arrests made by border patrol agents |
| 15 | Healthcare Provision | 1:55 | MSNBC News segment with panelist describing how healthcare is a human right and should be provided regardless of legal status in the country |

| | | | |
|---|---|---|---|
| 16 | Healthcare Provision | 1:32 | Fox News segment indicating that $2 billion of Medicaid funds is spent on providing emergency care to illegal immigrants, which is a practice that is in violation of existing laws. |
| 17 | Dream Act | 0:56 | CNN interview with Rep. Carlos Curbelo (R) on a Republican-led bill to provide a pathway to legal status for Dreamers |
| 18 | Dream Act | 0:57 | Segment from The Young Turks with panelist describing how the Dream Act encourages dangerous border crossings and puts children's lives at risk |
| 19 | Dream Act | 1:42 | Remarks by Senator Dick Durbin (D) describing and advocating for the Dream Act |
| 20 | Dream Act | 1:28 | Animated public service announcement on why the Dream Act is bad policy |
| 21 | Sanctuary Cities | 1:50 | AJ+ news segment describing sanctuary policies, including clips of gatherings in support of sanctuary cities and interviews with activists and undocumented immigrants |
| 22 | Sanctuary Cities | 1:30 | Fox News segment with panelist explaining why sanctuary policies are dangerous and illegal |
| 23 | Sanctuary Cities | 1:36 | Vice news segment showing clips of arrests made by Immigration and Customs Enforcement, discussing issues surrounding sanctuary policies, including how it lowers crimes. |
| 24 | Sanctuary Cities | 1:20 | Animated public service announcement describing the history of sanctuary cities and how the policies result in the release of 8000 convicted illegal immigrants. |
